# Structure and function of the intermembrane space domain of mammalian F_o_F_1_ ATP synthase

**DOI:** 10.1101/193045

**Authors:** Niels Fischer, Victoria Beilsten-Edmands, Dror S. Chorev, Florian Hauer, Chimari Jiko, Satoru Shimada, Kyoko Shinzawa-Itoh, Carol V. Robinson, Holger Stark, Christoph Gerle

**Affiliations:** Department of Structural Dynamics, Max Planck Institute for Biophysical Chemistry, Am Fassberg 11, Göttingen 37077, Germany; Department of Chemistry, Physical & Theoretical Chemistry Laboratory, University of Oxford, South Parks Road, Oxford, OX1 3TA UK; Institute for Protein Research, Osaka University, Suita, Osaka, Japan; Picobiology Institute, Graduate School of Life Science, University of Hyogo, Akoh, Hyogo, Japan; JST, CREST, Kawaguchi, Saitama, Japan

**Keywords:** *Bos taurus*, mitochondrial bioenergetics, membrane protein, mass spectrometry, OXPHOS

## Abstract

Mitochondrial F_o_F_1_ ATP synthase is a membrane bound molecular machine central to cellular energy conversion and cristae architecture. Recently, a novel domain has been visualized in the intermembrane space region of mammalian ATP synthase. The complete three-dimensional (3D) structure, composition and function of this domain - which we term intermembrane space domain (IMD) - are unknown. Here, we present two distinct 3D structures of monomeric bovine F_o_F_1_ ATP synthase by single particle cryo-electron microscopy (cryo-EM) that differ by the presence and absence of the IMD. Comparison of both structures reveals the IMD to be a bipartite and weakly associated domain of F_o_F_1_ ATP synthase. The tubular sub-domain of the IMD appears to contact the rotor-ring region, its globular sub-domain is anchored in the membrane-bending kink of the ATP synthase. However, absence of the IMD does not impact the kink in the transmembrane region ruling out a functional role in membrane bending. By combining our structural analysis with chemical cross-linking and reported biochemical, genetic and structural data we identify 6.8PL and DAPIT as the subunits forming the intermembrane space domain. We compare the present structure of the mammalian IMD in the bovine F_o_F_1_ ATP synthase monomer with structures of dimeric F_o_F_1_ ATP synthase from yeast and ciliate showing that the IMD is a common, but structurally divergent feature of several mitochondrial ATP synthases. On the basis of our analysis we discuss potential functions of the novel domain in rotary catalysis, oligomerization and mitochondrial permeability transition.

## Introduction

Mitochondrial F_o_F_1_ ATP synthase is central to the energy metabolism in mammalian cells producing more than 90% of ATP during respiration. To fulfill its energy transforming function, it utilizes the proton motive force (pmf) generated by the electron transport chain to recycle ATP from ADP+Pi via proton flow fueled rotary catalysis [1-6]. Moreover, mitochondrial ATP synthase plays an essential role in membrane bending and cristae formation [7-11]. Fragility and flexibility of this large membrane protein complex (>600 kDa, 17 different subunits) pose a major challenge for purification and structure determination of the entire complex. As a consequence, only crystal structures of stable sub-complexes are available, encompassing the membrane-extrinsic F_1_ domain, and the rotor-ring (c-ring) of the transmembrane F_o_ part [4, 12-15]. These structures provided important insights into the core function of ATP synthesis, while cryo-EM studies provided first lower-resolution structures of the intact complex [16-20].

Despite this progress, the precise localization of the mammalian-specific F_o_ subunits e, f, g, 6.8PL and DAPIT and their exact function is still unknown. The latter two subunits, 6.8PL and DAPIT, were only lately recognized as regular subunits due to their weak association with the complex [21-24]. Very recently, a novel domain on the intermembrane space side of mitochondrial F_o_F_1_ ATP synthase has been visualized in cryo-EM studies from diverse organisms: in the bovine monomeric ATP synthase complex [18, 19], as well as in the dimeric complexes of the yeast *Yarrowia lipolytica* [20] and the ciliate *Paramecium tetraurelia* [25]. The domain is only poorly defined in the bovine ATP synthase structures and the identity of the comprising subunits is unknown.

Here, we have used computational sorting of cryo-EM images to obtain a well-defined density for this novel domain of bovine F_o_F_1_ ATP synthase. We show that it is a separate structural domain – weakly associated with the remainder of F_o_ - which we term the intermembrane space domain (IMD). Based on the present structural and chemical cross-linking data and previously published studies, we deduce the bovine IMD to be formed by the subunits 6.8PL and DAPIT. We compare the structure and possible function of the mammalian IMD with the corresponding features of yeast and ciliate F_o_F_1_ ATP synthase.

## Results and Discussion

### Two compositional states of bovine F_o_F_1_ ATP synthase revealed by cryo-EM

In order to obtain a well-defined structure of the IMD of mammalian F_o_F_1_-ATP synthase we re-analysed our cryo-EM data of the monomeric *B. taurus* complex (Methods). In our first cryo-EM analysis [19] (Figure 1A left), as well as in the structures reported by the Rubinstein lab [18] (Figure EV2), the domain appeared as an elongated tubular feature of low density extending on the intermembrane side from the rotor-distal region of F_o_ close to the rotor. The very low density suggested strong structural heterogeneity due to flexibility or partial occupancy, impeding conclusive interpretation. Therefore, we now applied computational sorting of cryo-EM particle images to resolve the structural heterogeneity. We used two independent approaches to sort the data: i) supervised classification and ii) Maximum-likelihood based classification (Methods; Figures 1 and EV2). Both approaches yielded the same results; in the following, we will focus on the results obtained by supervised classification. In particular, we obtained two distinct structural states of bovine F_o_F_1_ ATP synthase – state 1 and 2 – each resolved at a resolution of about 12 Å (Figure 1A,B and Figure EV2). The difference density between the two states reveals no global variations, except an additional well-defined continuous density on the intermembrane space side in state 1, which is completely absent in state 2. The two states each encompass about 50% of the total particle population indicating that the poor definition of the IMD in the original map - obtained by averaging over all particles - was mainly the result of low occupancy.

**Figure 1.**
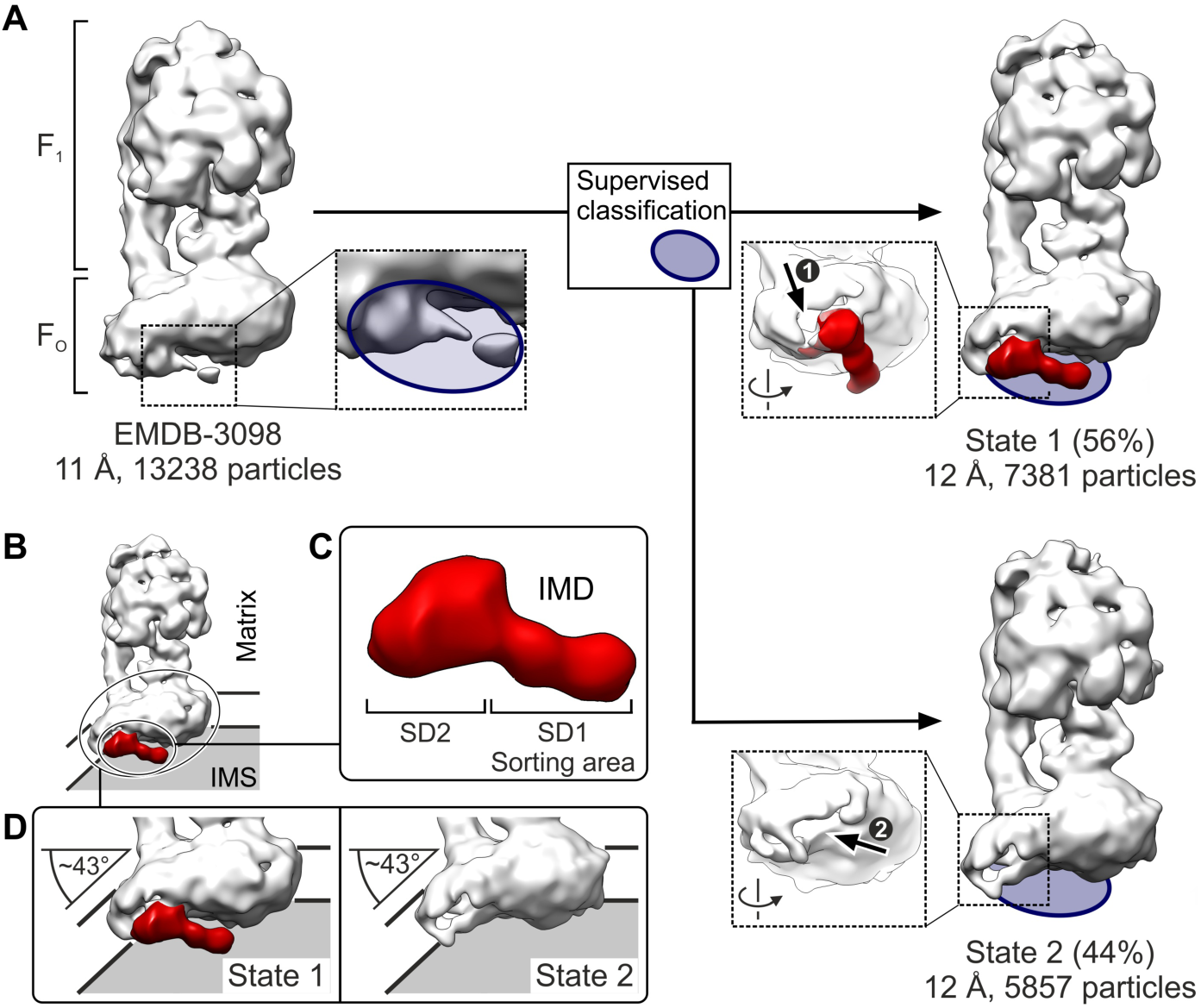
The intermembrane space domain (IMD) of mammalian F_o_F_1_ ATP synthase. A) Computational sorting of cryo-EM images reveals the IMD as a weakly associated domain of bovine F_o_F_1_ ATP synthase. Cryo-EM reconstructions rendered at ~3σ, as obtained from all particles [19] (top left) and from sub-groups after sorting (Methods, top and bottom right) with the resolution and number of particles noted for each state. Blue, density region used for sorting by supervised classification; red, difference density between the cryo-EM maps (state 1 minus state 2). Note the void between IMD and the tail of the F_o_ kink (arrow ❶) and the similar void in state 2 (arrow ❷). B) Location of the IMD. IMS, mitochondrial intermembrane space in grey. C) Structure of the IMD. SD1, tubular sub-domain; SD2, globular sub-domain. D) The IMD is not essential for the membrane bending kink of transmembrane F_o_. Comparison of the F_o_ kink in states 1 and 2, measured in degrees. The assumed position of the membrane is indicated by solid lines.

### Structure of the IMD of bovine F_o_F_1_ ATP synthase

The extra-density in state 1 reveals a more pronounced tubular sub-domain after computational sorting [18, 19] with an enclosed density that is sufficiently large to accommodate two alpha helices. It also includes a globular subdomain contacting the rotor-distal F_o_ region (Figure 1C). The complete absence of density for both sub-domains in state 2 strongly indicates that the sub-domains together form a stable domain, which is only weakly associated with the remainder of F_o_ via non-covalent interactions. Due to its location, we term this domain the intermembrane space domain, IMD.

Notably, the two states differ only with respect to the presence and absence of the IMD. In state 2 the IMD is completely absent, resulting in a half-pipe shaped opening on the rotor-distal side of the F_o_ region (Figure 1A bottom right). However, the membrane bending kink of the rotor-distal F_o_ region is identical between states 1 and 2 (Figure 1D). Consequently, the membrane bending function of mitochondrial bovine F_o_F_1_ ATP synthase is not directly linked to the IMD, despite the latter’s prominent size and position in the kink. Remarkably, there is also void in the IMD-bound structure, between the globular sub-domain of the IMD and the remainder of F_o_, which may stem from a water filled cavity or reflect the presence of bound lipids.

The tubular sub-domain of the IMD reaches towards the lipid plug protruding from the intermembrane space end of the rotor-ring. The structures of bovine ATP synthase reported by the Rubinstein lab similarly indicate a contact between the IMD and the rotor-ring region (Figure EV1C right). Whether the IMD interacts with subunits of the c_8_-ring or directly with the lipid plug protruding into the intermembrane space cannot be discerned at this stage.

### Subunit composition of the IMD

The exact localization of the bovine subunits e, f, g, 6.8PL and DAPIT in F_o_ is unknown to date (Figure 2A,B). To assign the subunit composition of the bovine IMD, as outlined below, i) we first performed biochemical and mass-spectrometric analyses of our purified complexes to verify their integrity and activity, ii) we surveyed the literature on the unassigned F_o_ subunits (see below and Table EV1), which consistently indicates 6.8PL and DAPIT to form the IMD and iii) we show that a model consisting of 6.8PL and DAPIT matches the present IMD structure and cross-linking data very well.

**Figure 2.**
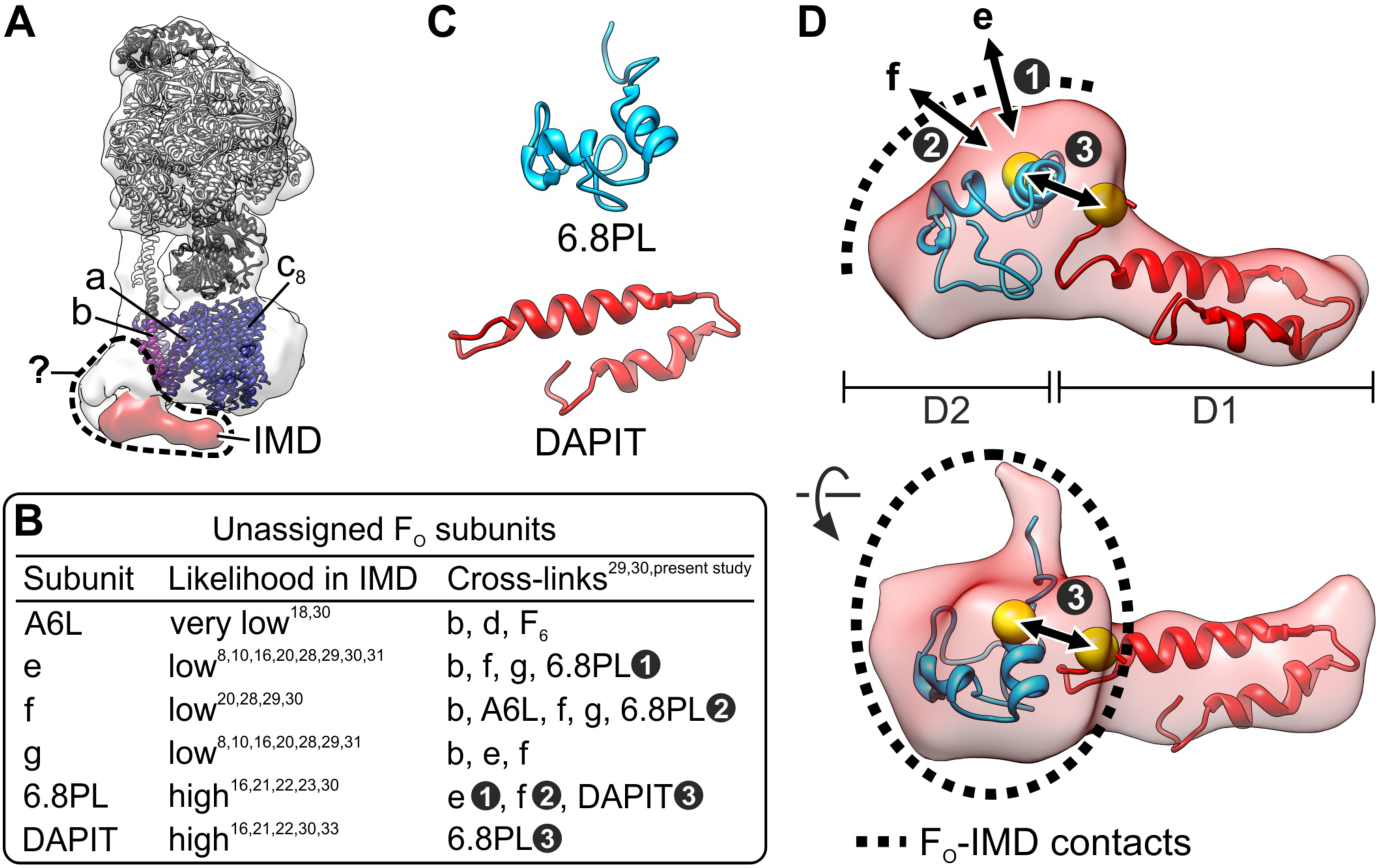
Structural identity of the mammalian IMD. A) Cryo-EM map of state 1 with fit of known structural elements of mammalian F_o_F_1_ ATP synthase (PDB-ID 5FIL) [18]. Grey, structures of F_1_ part; blue, c_8_-ring; purple, a-subunit; dark magenta, transmembrane part of b-subunit; dashed line, area of unknown subunit architecture. B) Unassigned subunits of F_o_ and their likelihood to be part of the IMD. The likelihood was deduced from reported data; see text and table EV1 for details. White numbers denote the cross-links in D). C) Homology models of F_o_ subunits 6.8PL (cyan) and DAPIT (red) obtained by iTasser [50]. D) Docking of subunits 6.8PL (cyan) and DAPIT (red) into the IMD density. The two-headed arrows denote previously reported cross-links between F_o_ subunits [30], the yellow spheres the C_α_ atoms of 6.8PL and DAPIT residues involved in cross-links. Dashed lines, contact area between the IMD and the remaining F_o_ part.

F_o_F_1_ ATP synthase was purified to homogeneity from bovine heart mitochondria by a combination of density gradient centrifugation and anion exchange chromatography in the presence of the novel, very mild detergent LMNG (also termed MNG-3) [26]. Analysis by denaturing SDS-PAGE and mass spectrometry demonstrated all known subunits of mitochondrial F_o_F_1_ ATP synthase to be present in the purified complex, in particular also the unassigned F_o_ subunits e, f, g, 6.8PL and DAPIT (Figure EV3). Furthermore, biochemical characterization by clear native PAGE and activity measurements indicated the complex to be intact, active and coupled.

Whereas the unassigned F_o_ subunits e, f, and g are clearly present in the purified complex, it is highly unlikely that these subunits are components of the IMD for several reasons (see also Table EV1). First, all three subunits have been shown to be tightly bound, integral components of the F_o_ part of bovine F_o_F_1_ ATP synthase [27]. This is exemplified by their persistent presence in F_o_F_1_ ATP synthase preparations, even when using relatively harsh purification procedures. For instance, all three subunits could be detected in the isolated F_o_ domain after stripping off the F_1_ domain and OSCP from submitochondrial particles by treatment with 3.3 M guanidine-HCL and subsequent solubilization with dodecyl-maltoside followed by mulitiple column chromatography [28]. This strong association of subunits e, f and g with the F_o_ region is incompatible with the low IMD occupancy (~50%) observed here under very mild purifications conditions. Second, cross-linking data have demonstrated that subunits e, f and g are arranged in close proximity to each other and to the peripheral stalk subunit b [29, 30] This proximity of e and g to b is further corroborated by detection of an assembly intermediate in human F_o_F_1_ ATP synthase comprising subunits b, e and g [31]. The peripheral stalk subunit b, in turn, is located distant from the IMD. Third, proteolysis experiments show the N-termini of subunits f and g to be exposed to the matrix side of the inner mitochondrial membrane [29], whereas the IMD is located on the opposite side. Furthermore, biochemical and structural data indicate that subunits e and g are indispensable for membrane bending of mitochondrial ATP synthase [8, 10, 32]. For example, the absence of subunits e and g in a recent cryo-EM study of monomeric yeast F_o_F_1_ ATP resulted in a structure lacking the membrane-bending F_o_ kink [25]. The present structural data (state 2) show that the IMD is not required for membrane bending. In line with our data, a ~40° kink in the rotor-distal domain of F_o_ was observed in a cryo-EM structure of bovine F_o_F_1_ ATP synthase, in which subunits e, f and g were present, but not 6.8PL and DAPIT [16].

The two subunits 6.8PL and DAPIT are the only subunits whose presence in the IMD is fully compatible with the reported data (Table EV1). Moreover, 6.8PL and DAPIT are stoichiometrically present in both monomeric and dimeric forms of the mammalian complex [21], but they are also known to easily detach from the F_o_F_1_ ATP synthase [21]. For instance, they are completely absent in preparations that lack sufficient amounts of phospholipids during purification with standard detergents such as dodecylmaltoside [33]. This behavior of 6.8PL and DAPIT very well correlates with the lower IMD occupancy obtained under very mild purification conditions here.

Integrating the data from crosslinking experiments, as well as genetic and structural studies, we therefore supposed that 6.8PL and DAPIT are the proteins forming the IMD in the bovine F_o_F_1_ ATP synthase. This prompted us to see whether models of the two subunits might fit our IMD structure. For each protein, we computed five homology models, which showed high consistency indicating good model quality (Methods, Figure 2C and Figure EV4). The best matching model of each protein was fit into the IMD structure, which suggested one distinct arrangement. Accordingly, DAPIT accounts for the elongated tubular sub-domain of the IMD, while 6.8PL occupies the globular sub-domain, anchoring the IMD to the remainder of F_o_ (Figure 2D). We then examined if this model is in line with our own and a recent cross-linking study [30] focusing on the unassigned bovine F_o_ subunits, including 6.8PL and DAPIT (Figure 3 and Figure EV5): i) the distance between lysine 49 of 6.8PL and lysine 55 of DAPIT perfectly matches the cross-linking data, ii) the positioning of DAPIT in the solvent-exposed tubular sub-domain of the IMD is in line with the lack of cross-links to other subunits and iii) cross-links of lysine 49 of 6.8PL with lysine 54 of subunit e and lysine 79 of subunit f corroborate the assignment of 6.8PL as anchor point of the IMD, which contacts the other F_o_ subunits. In addition, our own cross-linking experiments yielded cross-links connecting the c- and n-terminus of DAPIT (Figure 3 and Figure EV5), thereby confirming the computer generated hairpin structure. The high hydrophilic nature of the BS3 cross-linker and its efficacy in cross-linking DAPIT furthermore supports the notion that DAPIT resides within the intermembrane space. Interestingly, we also found that 6.8PL cross-links with the n-terminus of the c-subunit. Similar, as observed in cross-linking studies on the yeast complex, this cross-link may represent an inter-dimer cross-link, formed due to the presence of oligomeric F_o_F_1_ ATP synthase in our sample (Figure EV4).

**Figure 3.**
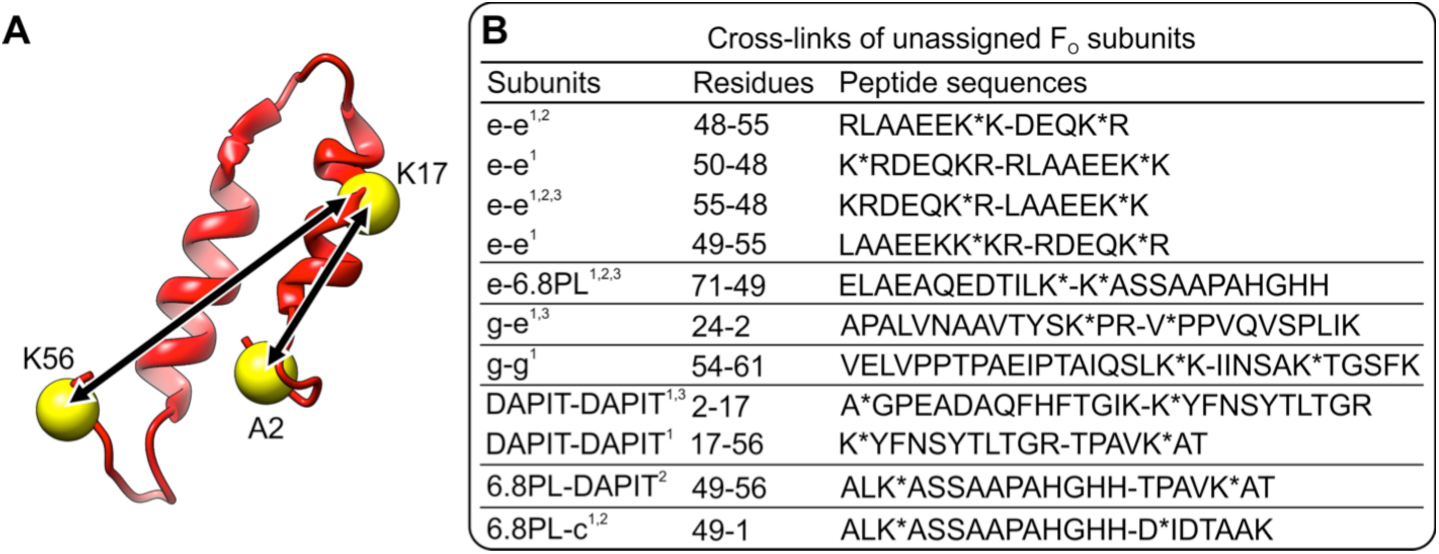
Cross-linking mass spectrometry is consistent with the modeled structure and intermembrane space position of DAPIT. A) Predicted structure of DAPIT with cross-linked residues indicated. B) Summary of cross-links identified with the crosslinkers BS3 (trypsin digestion) and DSS (trypsin/chymotrypsin or trypsin digestion) as indicated with 1, 2 and 3 in superscript, respectively. Note that all cross-links are in line with a position of DAPIT and 6.8PL on the intermembrane space side of the F_o_F_1_ ATP synthase.

In summary, our structural and chemical cross-linking data in combination with reported cross-linking, genetic and biochemical data strongly suggest that 6.8PL and DAPIT form the IMD, a loosely attached domain of mammalian F_o_F_1_ ATP synthase on the intermembrane space side.

### Structure and function of the IMD from mammalian vs. yeast and ciliate ATP synthase

A similar basic architecture for the IMD exists in diverse mitochondrial F_o_F_1_ ATP synthases, as evident from comparing the present bovine IMD structure with structures of dimeric F_o_F_1_ ATP synthases from the yeast *Y. lipolytica* [20] and the ciliate *P. tetraurelia* [25] (Figure 4). In all these ATP synthases, a structural feature – like the bovine IMD – extends on the intermembrane side from the rotor distal part of F_o_ towards the rotor ring suggesting that an IMD is a common feature of diverse mitochondrial ATP synthases. However, the shape of the IMD and its integration into the F_o_ region varies substantially in comparison to the bovine IMD. In yeast, the IMD is of similar shape and length, but more slender and seemingly more firmly integrated in the remainder of F_o_. In the ciliate enzyme, the IMD is very different in shape and substantially larger, likely more tightly associated with the F_o_ region due to a very large contact interface.

**Figure 4.**
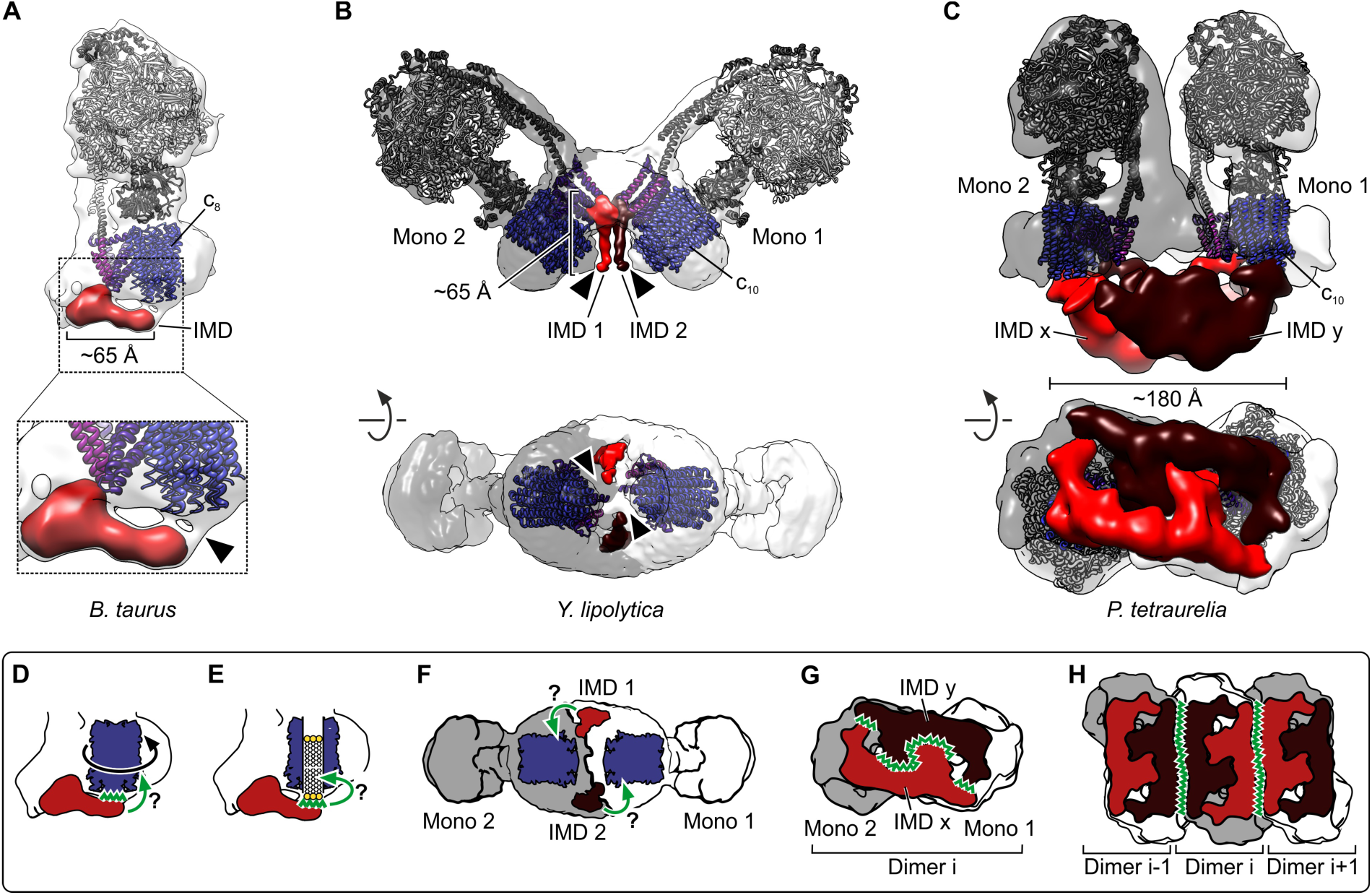
Structural and functional diversity of the IMD in mitochondrial F_o_F_1_ ATP synthases. Ribbon models are color-coded as in Figure 2a. A) Present structure of IMD-bound bovine ATP synthase (state 1), rendered at 3σ with fit of known elements (PDB-ID 5FIL) [18]. Bottom: Close-up rendered at 2σ indicating a potential interaction between the IMD and the rotor-ring region. B) Structure of dimeric yeast ATP synthase [EMDB-8152, PDB-IDs 5FL7 (F_1_-c_10_) and 5FIL] [18, 20]. The monomers (Mono 1 and 2) and their respective IMDs (IMD 1 and 2) are indicated. Note the close proximity between the IMD of one monomer and the rotor-ring region of the other monomer (arrow-heads), which contact each other at lower threshold (2σ, not shown). C) Structure of dimeric ciliate ATP synthase [EMDB-3441] with atomic models from yeast PDB-IDs 5FL7 (F_1_-c_10_) and Bos taurus 5FIL fitted for orientation [18, 20]. The monomer assignment of IMDs (IMD x and y) is unclear at present. Note the very large contact areas between the IMDs and the F_o_ rotor-ring regions. D-E) Function of mitochondrial IMDs. Potential functions are marked by green arrows and question marks, zig-zag lines mark interactions. D) Potential interference with rotary catalysis upon contact with the rotor-ring region. E) Potential functions in assembly or gating of the mitochondrial permeability transition pore upon contact with the lipid plug of the rotor-ring. F) Potential function in intra-dimer communication. G) Intra-dimer stabilization. H) Inter-dimer contacts specific for formation of helical ATP synthase arrays in *P. tetraurelia* [25].

Notably, the IMD appears to contact the F_o_ rotor-ring region in all three cases. This preserved contact may play a role in regulating this functionally important region (Figure 4D and E). The exact contact sites on the rotor-ring, i.e. the c-subunits and/or the lipid plug, are not clear at present. A direct contact with the rotor-ring c-subunits would interfere with rotary catalysis. In this case, the IMD could function as an inhibitor of F_o_ - in analogy to the inhibition of the F_1_ region by IF1 [34] - by binding to the c-subunits. Alternatively, a contact of the IMD with the lipid plug protruding from the lumen of the rotor-ring would not impede rotary catalysis. Such a contact might help in stabilizing the F_o_ structure in accordance with the suggested role of 6.8PL and DAPIT as assembly factors [35, 36].

Furthermore, a contact with the lipid plug might allow manipulating the lipids filling the rotor-ring lumen. Such a scenario would be of particular importance in the mammalian system, as Bernardi and colleagues recently proposed a completely novel role for mammalian F_o_F_1_ ATP synthase as the mitochondrial permeability transition pore [37]. Accordingly, the mammalian IMD might be involved in gating of the mitochondrial permeability pore by controlling the lipid plug [38] or the dimer interface. A function of the rotor-ring lumen as the pore is currently a matter of debate [39, 40].

In the dimeric ATPases from yeast and ciliate, the IMD of one F_o_F_1_ monomer interacts with the neighboring monomer. These intra-dimer contacts could principally serve two different roles. First, such contacts might facilitate intra-dimer communication: In the yeast dimer the IMD of one monomer contacts the rotor-ring of the other monomer, which may provide a means for allosteric intra-dimer regulation (Figures 4B, C, F). Second, intra-dimer contacts might stabilize dimerization, as suggested by the very large, entangled interaction interface between the IMDs in the ciliate dimer (Figure 4C, G). Furthermore, in *P. tetraurelia* the IMDs make extensive inter-dimer interactions (Figure 4H) that promote the formation of helical synthase oligomers, which, in turn, may explain the tubular cristae specific for this ciliate [25]. A structure of the mammalian ATP synthase dimer will be required to reveal whether also the mammalian IMD exerts dimer-specific functions.

In conclusion, the present data show that bovine F_o_F_1_ ATP synthase can exist in two compositional states, which differ by the presence or absence of the IMD – a weakly associated domain on the intermembrane space side of the F_o_ region. The IMD-bound structure in combination with the cross-linking and previous data strongly suggests that the two hitherto unassigned F_o_ subunits 6.8PL and DAPIT form the IMD. Accordingly, 6.8PL constitutes the globular subunit of the IMD, which is associated to the rotor-distal F_o_ region. Despite its prominent position at the membrane bending F_o_ kink, our data clearly show that the IMD is not essential for kink formation. DAPIT forms the tubular sub-domain of the bovine IMD, which extends towards the rotor-region, suggesting a potential function of the IMD in regulation of this functionally important F_o_ part. Although present in F_o_F_1_ ATP synthase complexes from evolutionary divergent organisms, the observation of IMD-like features seems to be limited to the mitochondrial enzyme [41]. Moreover, the absence of an IMD in dimers isolated from the colorless green alga Polytomella [17] indicates that an IMD may not be present in all mitochondrial F_o_F_1_ ATP synthases. The present work reveals novel insights into the structure and function of the IMD providing the basis for future approaches towards an understanding of dimer stabilization, mitochondrial chemiosmosis and apoptosis.

## Materials and methods

### Purification of intact, active and coupled bovine F_o_F_1_ ATP synthase using LMNG

Protein purification was conducted as previously described for the 2D crystallization of bovine F_o_F_1_ ATP synthase [11, 42] including modifications for LMNG stabilized complexes as described for the GraDeR procedure [19, 43]. Briefly, fresh bovine hearts were obtained immediately after slaughter and inner mitochondrial membranes were purified as previously reported [44]. Sub-mitochondrial particles were kept in in 40 mM HEPES pH 7.8, 2 mM MgCl2, 0.1 mM EDTA, and 0.1 mM DTT and solubilized on ice via addition of deoxycholate and decylmaltoside to final concentrations of 0.7% (wt/vol) and 0.4% (wt/vol), respectively. Subsequently, the suspension was centrifuged at 176,000×g for 50 min and the supernatant applied to a sucrose step gradient (40 mM HEPES pH 7.8, 0.1 mM EDTA, 0.1 mM DTT, 0.2% [wt/vol] decylmaltoside and 2.0 M, 1.1 M, 1.0 M, or 0.9 M sucrose) and centrifuged at 176,000×g for 15.5 hr. Fractions exhibiting ATPase activity determined by an ATP-regenerating enzyme-coupled assay [45] were loaded onto a Poros-20HQ ion-exchange column. The detergent was exchanged to LMNG using a double gradient from 0.2% to 0% decyl-maltoside and 0%–0.05% LMNG for 80 min at 1 ml/min. Complexes were eluted by a linear concentration gradient of 0–240 mM KCl in 40 mM HEPES pH 7.8, 150 mM sucrose, 2 mM MgCl2, 0.1 mM EDTA, 0.1 mM DTT and 0.05% (wt/vol) LMNG. Shortly after elution, ATP synthase fractions containing high amounts of native phospholipids as determined by ammonium molybdate complexation were flash frozen in aliquots of about 500 μl for later use.

### Gel electrophoresis, ATPase activity measurement and mass spectrometry

To confirm the intactness of the F_o_F_1_ ATP synthase in the sample used for cryo-grid preparation aliquots of the same purification were subjected to non-denaturing clear-native PAGE [46]. The specific enzymatic activity and the proportion of coupled complexes were verified by an ATP-regenerating enzyme-coupled assay [45]. The subunit composition was examined by denaturing SDS-PAGE and the presence of the lower molecular weight subunits (<15,000 Da) was confirmed by MALDI-TOF mass spectrometry (Bruker Daltonics Inc., MA, USA).

### Cryo-EM analysis

As previously reported, we used the GraDeR approach for cryo-EM preparation of the purified ATP synthase complexes, which allowed us to determine an 11 Å cryo-EM structure of the bovine F_o_F_1_ synthase complex by averaging over 13,000 cryo-EM particle images [19]. The scattered density observed for a novel, tubular structural feature on the intermembrane side of the complex prompted us to re-analyse our cryo-EM data set here. We used two independent approaches – supervised classification and Maximum-Likelihood based (ML) classification – to sort the particle images computationally and to improve visualization of this novel feature.

Supervised classification (Figure 1A) – We first re-refined the set of 13,000 cryo-EM particle images using a larger mask encompassing the tubular feature. The resulting map showed only a slightly improved density for the tubular feature. Automated segmentation of this map in UCSF chimera indicated that the tubular feature is connected to a globular region in the rotor-distal kink of F_o_, both together potentially forming a domain of F_o_. Therefore, we devised a mask encompassing this domain to create two versions of the cryo-EM map one with and one without this domain with the software UCSF Chimera [47]; to avoid high-resolution reference bias the maps were low-pass filtered to 18 Å. These maps were then used to sort the particle images by supervised classification resulting in two sub-groups (state 1: 7381 particles; state 2: 5857 particles). Supervised classification was performed without alignment, i.e. the particles were aligned first against the average map and then sorted according to their best match with one of the two map versions. For each sub-group, a 3D reconstruction was computed by gold-standard refinement in Relion [48], amplitude-sharpened and low-pass filtered to the respective final resolution (12 Å, Figure EV3A). In comparison to the reported 11 Å map computed from all particles, the structure of state 1 shows more prominent and better defined density for the tubular feature and for the globular region. The structure of state 2, in contrast, displays no density for the tubular feature and also misses the globular region, whereas the density of the remaining F_o_ kink region also improved.

ML classification (Figure EV2A) – The 13,000 cryo-EM particle images were binned to a pixel size of 5.61 Å/pixel to improve speed and signal-to-noise ratio for subsequent ML-classification. Focused 3D ML-classification in Relion was applied to sort the binned data into two sub-groups according to the major structural differences variations in the F_o_-region. We used a feature-less elliptical mask entailing the entire F_o_ region to avoid any bias in the classification. For each of the two resulting sub-groups, a 3D structure was computed from the corresponding not-binned particle images, amplitude sharpened and low-pass filtered to the final resolution: State ML-1 was computed from 8131 particles at 12 Å resolution, state ML-2 from 5107 particles at 13 Å resolution.

Image processing was performed using Imagic-5 [49], Relion 1.3 and Relion 2.0 [48]. Homology modelling of subunits 6.8PL (Uniprot P14790) and DAPIT (Uniprot Q3ZBI7) of bovine F_o_F_1_ ATP synthase was performed using iTasser [50]. Rigid-body fitting of known structural elements was performed using UCSF Chimera [47]. Fitting of models indicated that the original pixel size (1.6 Å/pixel) [19] was inexact; accordingly, the pixel size was corrected to 1.53 Å/pixel. To model the present cryo-EM maps the best matching structure (PDB-ID 5FIL) was chosen from the available models for different sub-states of bovine ATP synthase [18]. Segmentation of cryo-EM maps was performed using the SEGGER tool [51] implemented in UCSF Chimera [47].

### Chemical cross-linking

6 μL ATP synthase complex was cross-linked with either BS3 cross-linker (1:1 ratio BS3-d0:BS3-d4) at a final BS3 concentration 0.6 mM or DSS cross-linker (1:1 ratio DSS-d0:DSS-d12) at a final DSS concentration 0.7 mM at 26^o^C for 1 hour in a thermomixer (Eppendorf) at 650 rpm. The cross-linking reaction was quenched for 15 minutes at room temperature in a thermomixer at 650 rpm. The samples were separated by SDS-PAGE and protein bands excised and digested as described [52] using trypsin for BS3 samples and both chymotrypsin and trypsin for DSS samples.

Peptides were separated on an Ultimate 3000 UHPLC system (Thermo Fischer Scientific). The peptides were trapped on a C18 PepMap100 pre-column (300 μm i.d. x 5 mm, 100Å, Thermo Fisher Scientific) using solvent A (0.1% formic acid) at a pressure of 500 bar. The peptides were separated on an in-house packed analytical column (75 μm i.d. packed with ReproSil-Pur 120 C18-AQ, 1.9 μm, 120 Å, Dr.Maisch GmbH) with a gradient of 15-38% (vol/vol) mobile phase B (100% ACN/0.1% formic acid) over 35 min then 38-58% mobile phase B for 15 min, with a column temperature of 45^o^C. The peptides were eluted into a QExactive Hybrid Quadrupole-Orbitrap mass spectrometer (Thermo Fischer Scientific) via an EASY-Spray nano-electrospray ion source (Thermo Fischer Scientific) for analysis. The mass spectrometric conditions were: spray voltage 2.1 kV; capillary temperature 320 °C. The Orbitrap was operated in data-dependent mode. Spectra were acquired in the orbitrap (*m/z* 350-1500) with a resolution of 70,000 and an automatic gain control (AGC) target at 3 × 106. The ten most intense ions were selected for HCD fragmentation in the orbitrap at an AGC target of 50,000. Singly and doubly charged ions and ions with unknown charge states were excluded from the analysis.

Raw data were converted to mgf format with pXtract [53] and cross-linked peptides identified with pLink software [54]. pLink search parameters included the following: instrument HCD; variable modification Oxidation [M]; fixed modification Carbamidomethyl [C]; max. missed cleavage sites 3. Spectra of cross-linked peptides were validated by manual inspection.

### Accession numbers

The cryo-EM maps of states 1 (IMD-bound) and 2 (without IMD) of mammalian F_o_F_1_ ATP synthase have been deposited in the Electron Microscopy Data Bank with accession codes EMD-YYYY and EMD-ZZZZ, respectively. Cryo-EM micrographs and particle images have been deposited in the EMPIAR data bank with accession code EMPIAR-ZZZZ.

## Acknowledgements

We would like to thank Tomitake Tsukihara for his advice and support during the whole project, Shinya Yoshikawa for his support in establishing purification procedures and Bernhard Ludewig for the graphical abstract. This work was supported by the JST, CREST Grant JPMJCR12M3 (to Tomitake Tsukihara and CG), the Platform for Drug Design, Discovery and Development from MEXT, Japan (to CG), the Grants-in-Aid for Scientific Research (Challenging Exploratory Research: 15K14464) from MEXT, Japan (to CG), the Grants-in-Aid for Scientific Research (Kiban B: 17H03647) from MEXT, Japan (to CG), the JSPS Grant 25-05370 (to CJ)), the European Research Council grant number 69551-ENABLE (to CVR) and the Max Planck Society (NF, FH and HS). VB-E acknowledges funding from the Engineering and Physical Sciences Research Council and DSC acknowledges funding from the European Research Council grant ENABLE.

## Author contributions

NF performed the cryo-EM analysis, VB-E and DSC performed the cross-linking and mass-spectrometry analysis, CVR supervised mass-spectrometry, CJ, SS and KS-I performed protein purification and biochemical experiments, NF and CG conceived the project, interpreted the data and wrote the paper with inputs from all authors.

## Conflict of interest

The authors declare that they have no conflict of interest.

## Figures

**Figure EV1.**
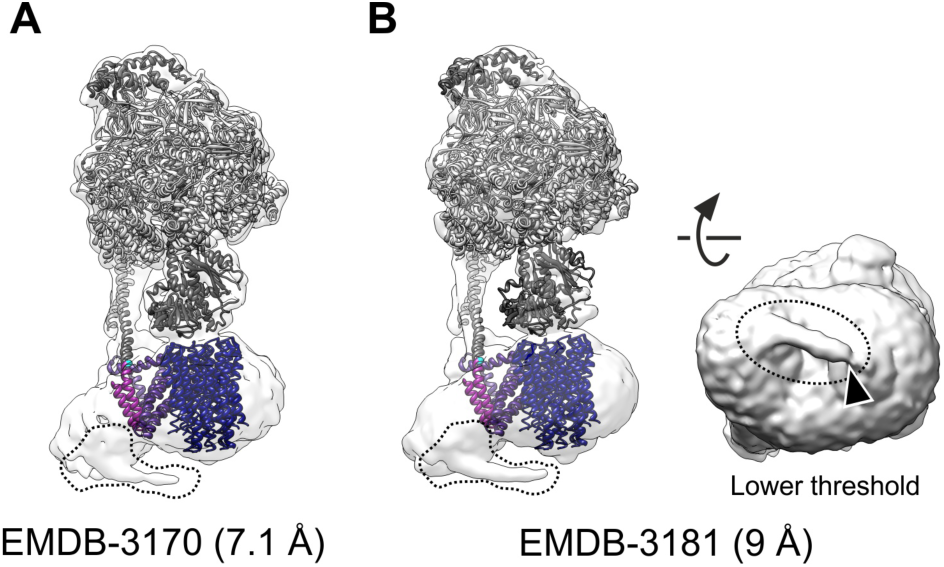
IMD density in cryo-EM structures of bovine F_o_F_1_-ATP synthase. The dashed line indicates the contour of the IMD from the present study. Ribbon models of subunits (PDB-ID 5FIL) [18] are color-coded as in Figure 2a. A) Structure of state 3b depicting typical density for the IMD as observed similarly in the other states reported in [18]. B) Average structure from all states obtained by aligning only with respect to the F_o_ region, resulting in a slightly better defined density for the IMD [18]. Right: View from intermembrane space side rendered at lower threshold indicating an interaction between the IMD and the rotor-ring of F_o_.

**Figure EV2.**
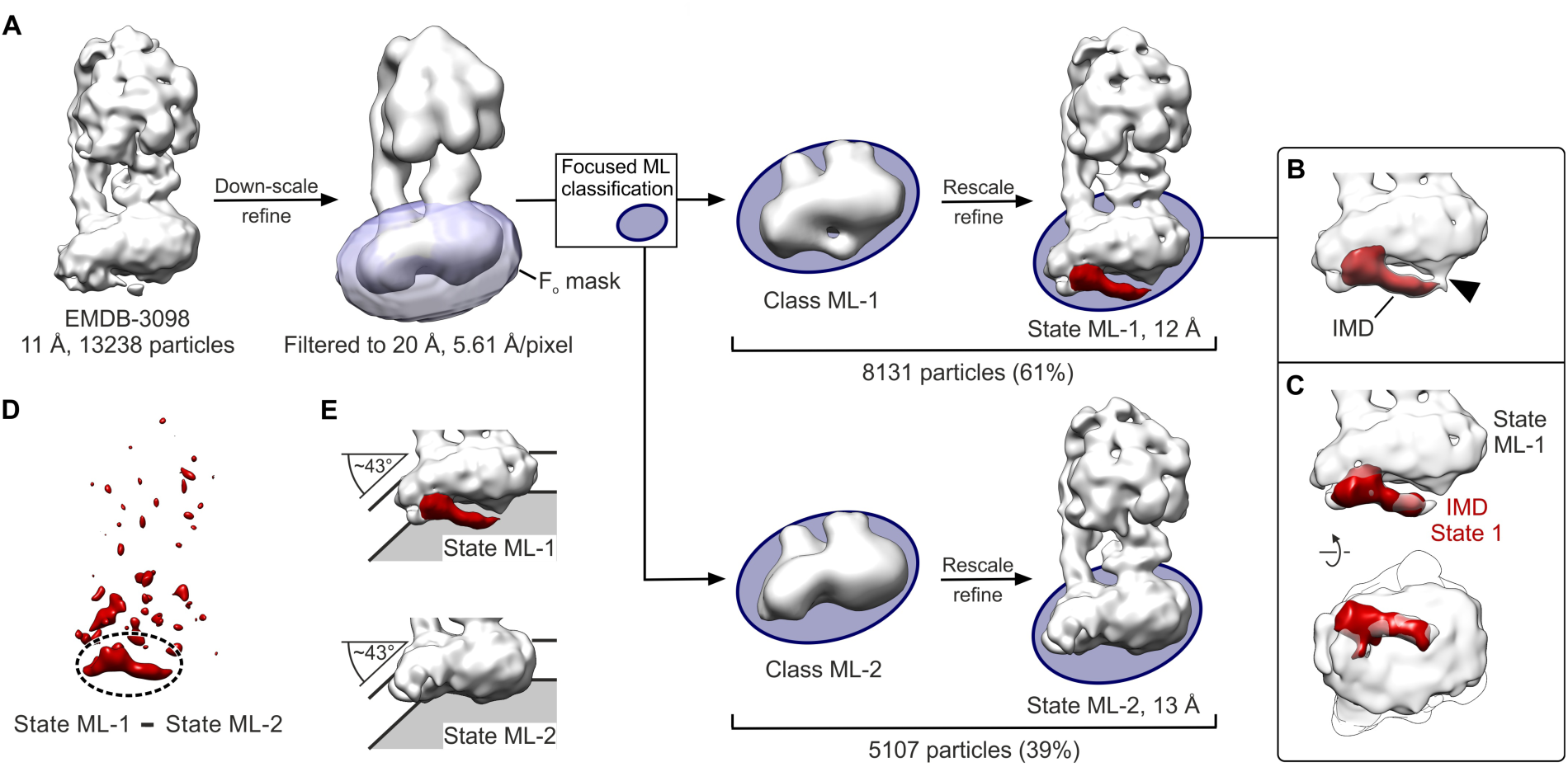
Maximum Likelihood-based (ML) classification of cryo-EM images confirms the results from supervised classification. A) ML classification of particle images focused on the F_o_ region. Cryo-EM maps are shown at ~3 σ, as computed from i) all particles (left), ii) all particles at smaller pixel size (second from left), iii) sub-groups in the last iteration of classification masked to the F_o_ region (second column from right) and iv) sub-groups after sorting at the original pixel size with the respective resolution in Å. Purple, mask used for classification focused on the F_o_-region; red, density for the IMD. B) Close-up of the F_o_ region and the IMD (red) in state ML-1. Note the contact between IMD and rotor-ring at slightly lower threshold (2.5 σ). C) Similarity of the bipartite IMD structure as obtained by different sorting approaches. Transparent white, density for F_o_-region as obtained by ML classification; red, IMD density as obtained by supervised classification. D) Difference density between states ML-1 and ML-2 identifies weak association of the IMD as the major variation in the F_o_-region. Dashed circle, difference density corresponding to the IMD. E) Similar membrane bending kink of F_o_ in the presence and absence of the IMD as observed in the sub-groups from ML classification.

**Figure EV3.**
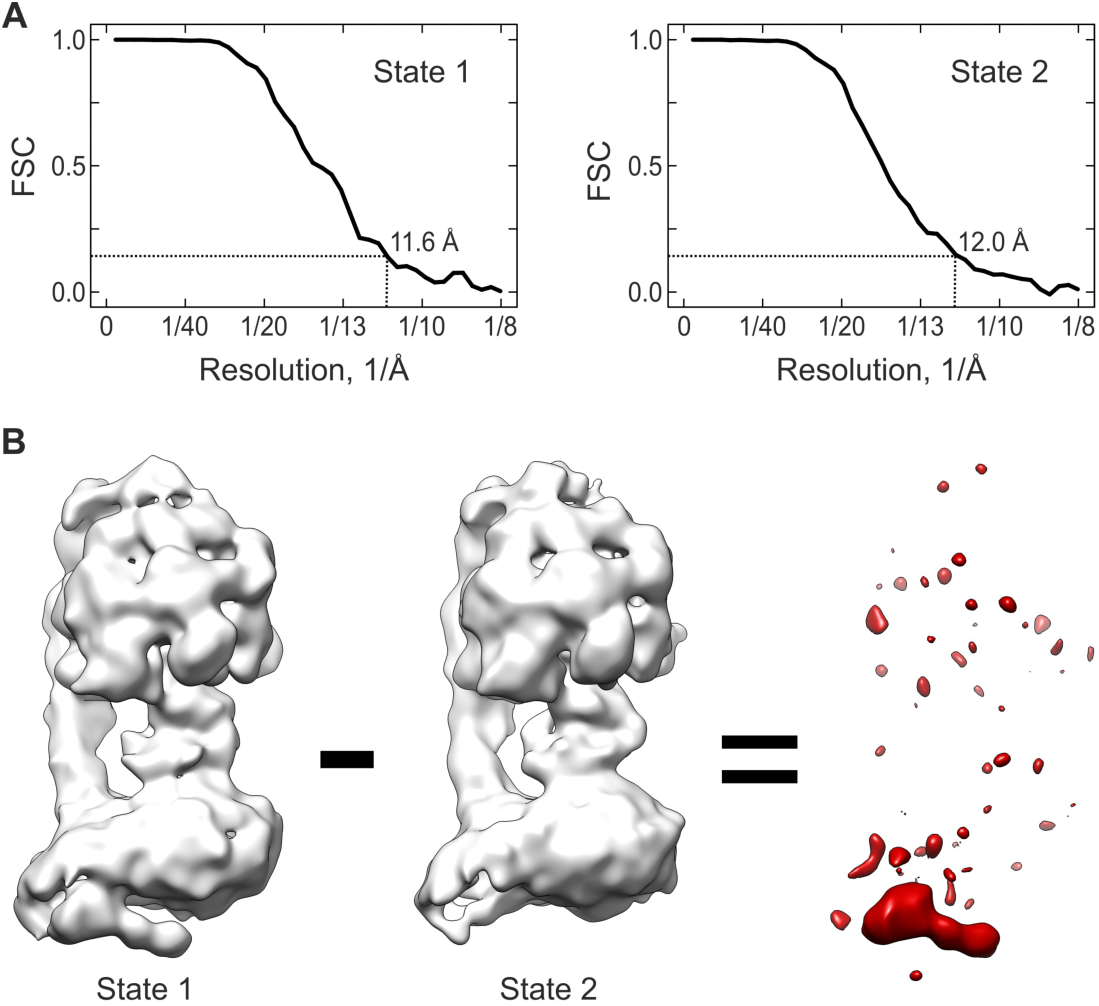
Cryo-EM analysis. A) Fourier-Shell-Correlation (FSC) curves for state 1 (left) and 2 (right). The vertical black dashed lines indicate the resolution according to the 0.143 criterion. B) Difference density (red) as computed between states 1 and 2.

**Figure EV4.**
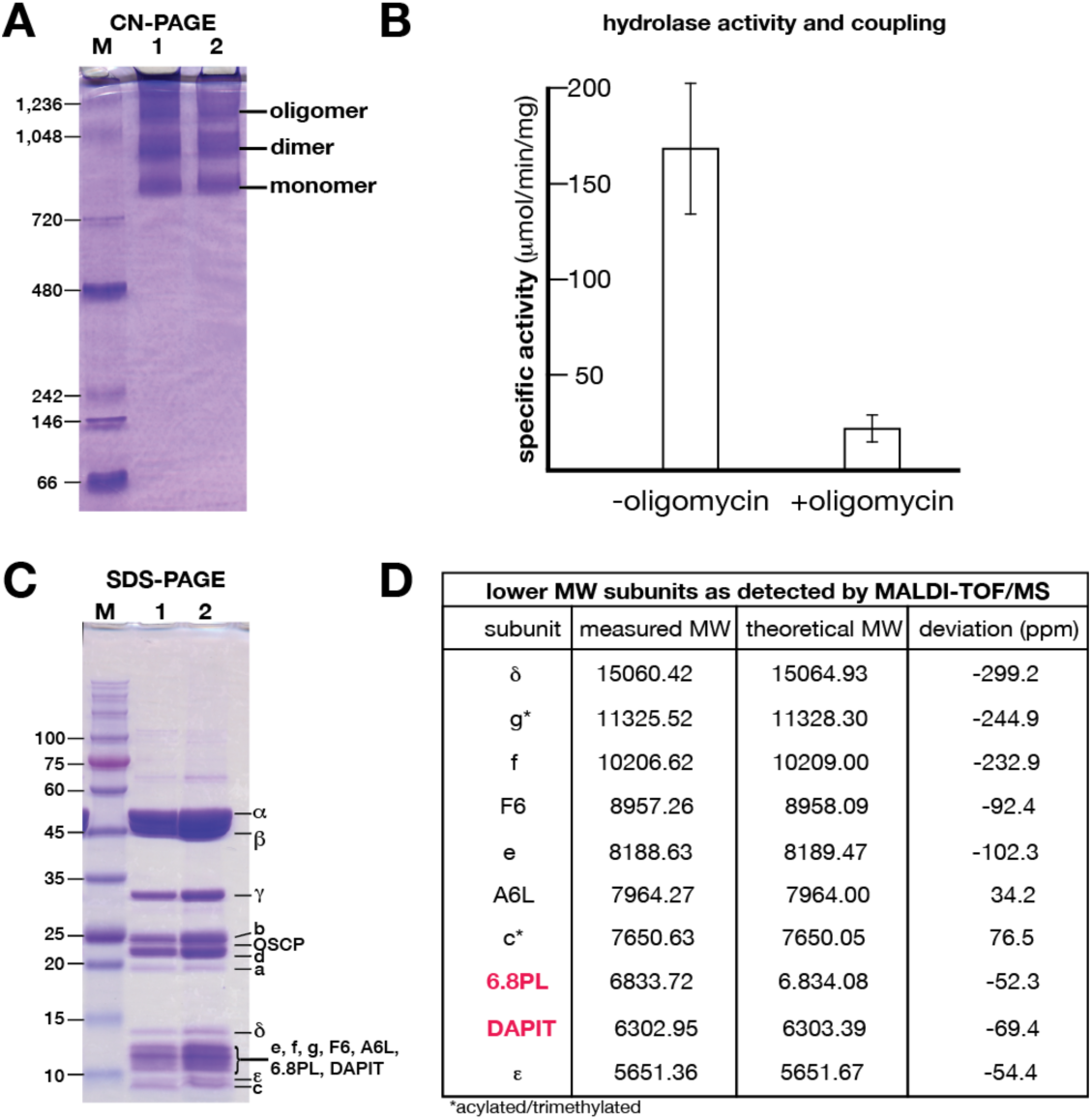
Integrity and activity of LMNG-purified bovine F_o_F_1_ ATP synthase complexes. A) Oligomeric states as observed by clear native PAGE (4-16%) gel electrophoresis [46]. Lane M: molecular weight markers as indicated; Lane 1 & 2: F_o_F_1_ ATP synthase (30 μg). B) Hydrolase activity and coupling as measured by an NADH oxidation coupled enzymatic assay [45]. Error bars indicate the standard error from three independent purifications. C) Subunit composition as determined by SDS-PAGE (10-20%) gel electrophoresis. Lane M: molecular weight markers as indicated; Lane 1 & 2: F_o_F_1_ ATP synthase (30 μg). D) Lower-weight subunit composition by mass-spectrometry. All expected subunits in the range up to 15 kDa were detected, including the very weakly associated subunits 6.8PL and DAPIT (red).

**Figure EV5.**
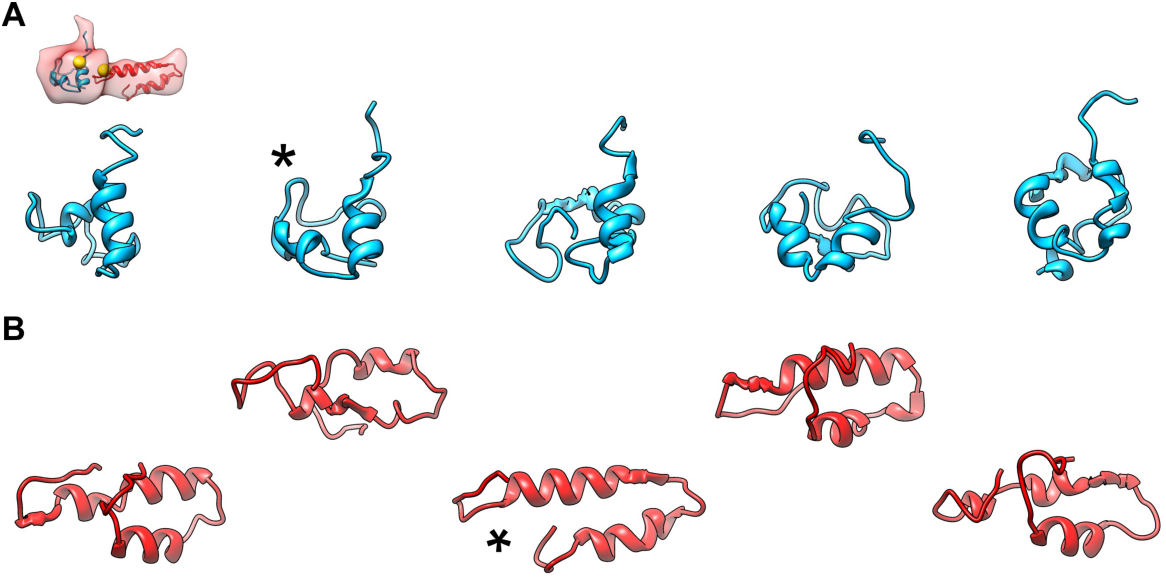
Homology models of 6.8PL and DAPIT as obtained by iTasser. Asterisks indicate the models used for fitting. A) Structural models of 6.8PL. B) Structural models of DAPIT. Note the two-helix bundle preserved in all five models matching the tubular sub-domain of the IMD.

**Figure EV6.**
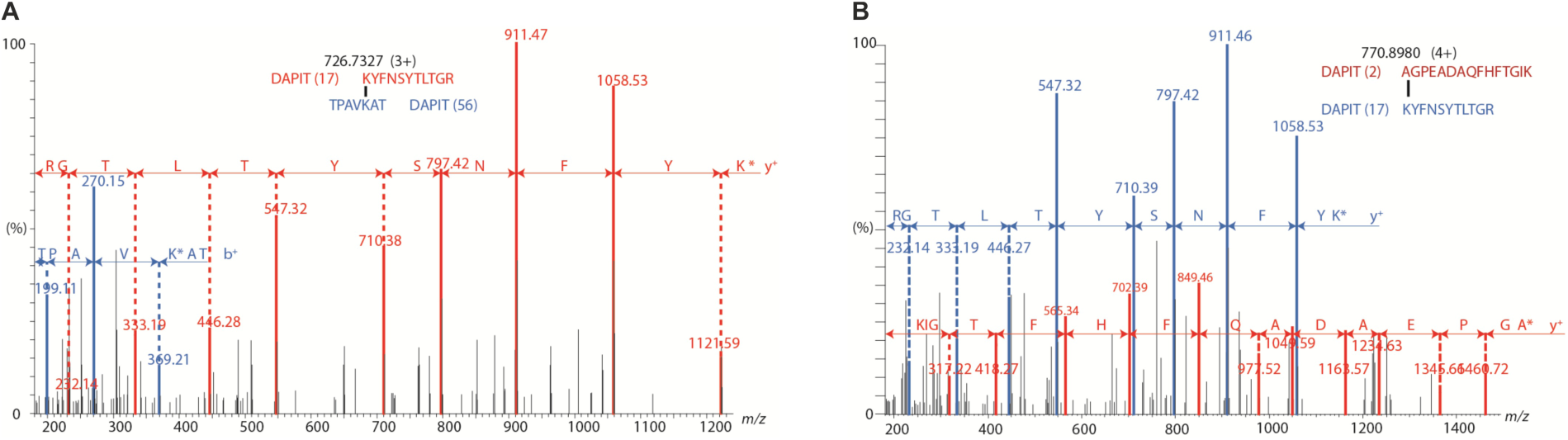
Tandem MS analysis of DAPIT peptides cross-linked by BS3 viaK17- (A) and A2-K17 (B).

**Table EV1.**
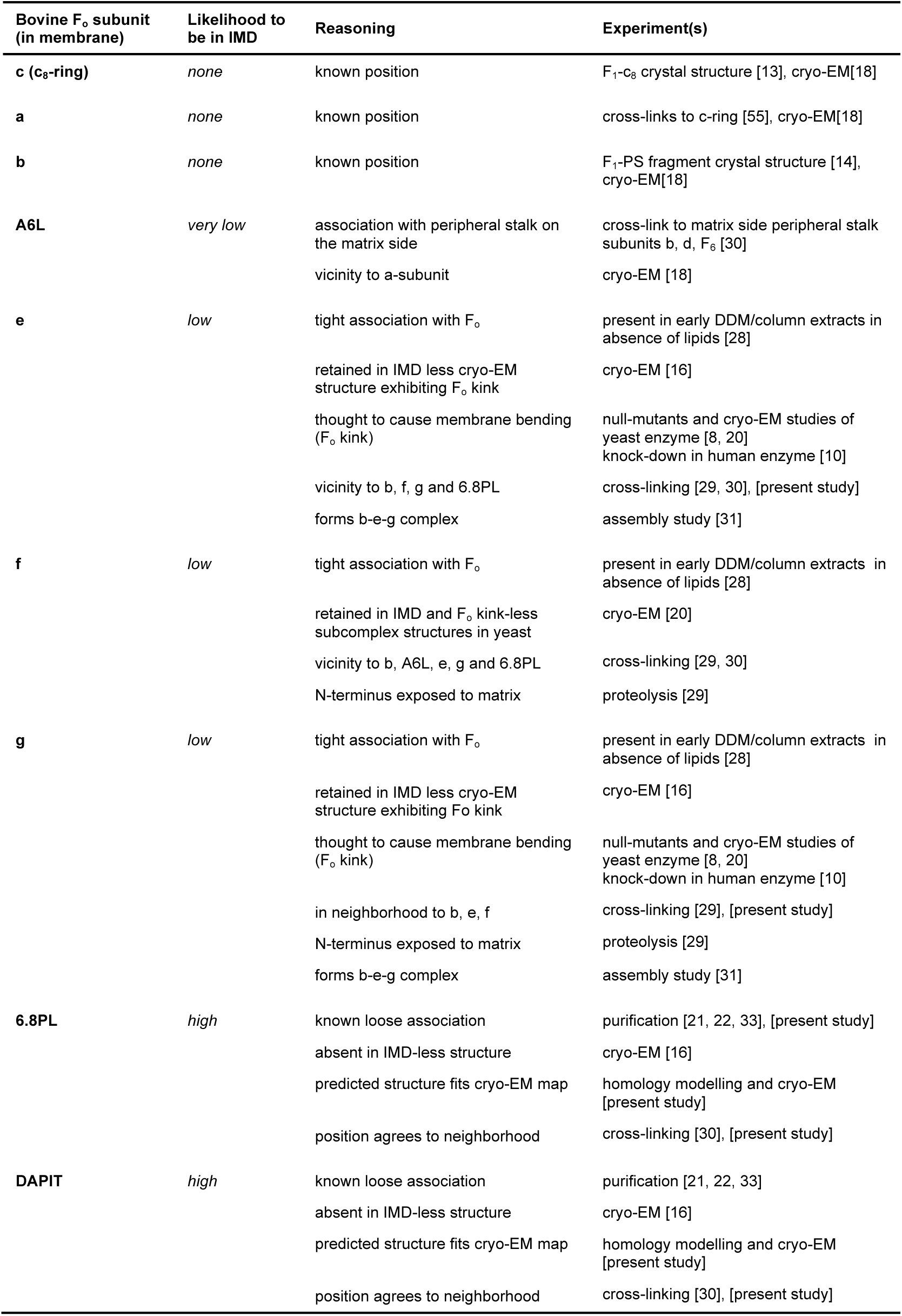
Identity of the intermembrane space domain (IMD) in the light of reported experimental data.

